# Multispecies relations shape bird-feeding practices

**DOI:** 10.1101/2024.12.19.629318

**Authors:** Tuomas Aivelo, Mikko Aulio, Johanna Enström, Purabi Deshpande, Anna Haukka, Heta Lähdesmäki, Katja Rönkä, Andrea Santangeli, Virpi Väkkärä, Aleksi Lehikoinen, Rose Thorogood, Anttoni Kervinen

## Abstract

While humans often feed birds in their backyards, there is growing awareness that this has positive and negative effects on local biodiversity. Whether the observed species assemblage shapes human activities has, however, rarely been investigated. We analyzed 15,088 open-ended answers from 9,473 Finnish respondents about why they have increased or reduced feeding birds. They mentioned 58 avian and non-avian species linked to changed practices. The main reasons for change were 1) respondent’s relation to non-human species, 2) respondent’s relation to other humans and 3) relations between non-human species. Most taxa and reasons could lead to both increase or decrease in feeding, although the direction was context-dependent. We suggest that bird-feeding is an interactive process where the species community strongly affects feeding practices, which in turn can affect community composition. Recognising this process is crucial for understanding the effects of bird-feeding on both humans and nature and providing more nuanced guidance.

## Introduction

Intended (and unintended) provisioning of wildlife is widespread throughout the world. Across human cultures, people feed animals for different reasons, such as hunting, maintaining ecosystem services or performing scientific studies, but it is especially common as a hobby^1^. Indeed, bird-feeding might be the most common interaction between humans and wild animals in the Global North^2^ and constitutes one of the most common and consequential forms of care that humans give towards other species. While feeding birds brings benefits to people, there is nevertheless growing concern that bird-feeding practices may have unintended consequences on nature^3^. For example, in the UK, three times more feed intended for birds is sold annually than the populations of the ten most common garden birds can theoretically consume^4^. While providing such food enables some species to better survive the winter or increase breeding productivity, it can also shape synanthropic bird^5,6^ and non-avian populations^7^ and have wide-ranging effects on ecological communities through pathogens^8^, competition^9^ or predation^10,11^. All in all, it is still unclear whether the overall effect of bird-feeding on biodiversity is net positive or negative^4,12^. As bird-feeding is a culturally and socially varying practice (such as when, where, by whom and how birds are fed)^6^, if we are to understand how bird-feeding affects ecological cascades and biodiversity, it will be crucial to elucidate the drives behind changes in these human practices.

The essential driver of bird-feeding is simply that people want, for various reasons, to feed birds^6,12^ In multiple studies of their motivations, people report that feeding aesthetically pleasing species, or caring for species perceived as in need, brings pleasure and enhances their own wellbeing^2,13,14^. This may come from a desire to nurture nature, to enhance opportunities to observe and educate themselves about different species, to provide a sense of companionship, or even to make amends for apparent negative effects on nature^13^. Bird-feeding is also increasingly framed as a tool to “(re)connect” people with nature^15,16^, echoing the pervasive Western human-nature divide^17^. However, and perhaps due to this emphasis on benefits to people, there are still relatively few studies on how the species communities that are attracted or do not come to bird-feeding sites, whether targeted or not, affects humans’ bird-feeding practices ^1,6,12^.

Bird-feeding has a broad non-human component that could influence decisions of what, where, and when to feed. In one of the few studies investigating how observations of nature influence feeding activities, Dayer at al. ^12^ found that observing a predator such as a domestic cat was coupled with a range of actions: for example, respondents scared off the cat, improved the shelter available for birds, and relocated feeders to make them less accessible to cats. People may also adjust their behaviour differently depending on variation how they value the species they observe, as there is ample evidence that people value different species in different ways^18^. Dayer et al.^12^ found that observing a hawk at feeders led some respondents to make similar changes as in relation to the presence of cats, but in contrast, half of the respondents stated that a hawk would not change any of their feeding practices due to it being a ‘natural predator’ and part of the ‘circle of life’. Even perceived interactions with ‘undesirable’ non-target species can potentially have large impacts on changes in bird-feeding actions. In a recent study using part of the dataset that we use here ^19^, the risk of rats being attracted to the site was the most common reason given for why people had reduced their bird-feeding activities – and this seemed to be encouraged more by rules and regulations rather than direct observation of rats. The classification between wanted and unwanted species is also manifested in the apparatus used to provide food. Indeed, the bird feeder can be seen more of a technology of exclusion as avoiding feeding unwanted species drives the evolution of the bird feeder models more than improvements for feeding the species preferred^20^. Bird-feeding can therefore be considered an interesting case of human value judgements and trade-offs between competing interests, and should be studied as a continuous multispecies interaction where humans offer food, non-human species either take up or do not take up the offer, and humans adjust their feeding practices accordingly^20^.

In this study, we therefore focus on multispecies relations to look at the drivers that lead to changes in bird-feeding practices. There is ample time for people’s preferences and multispecies interactions to manifest, as bird-feeding is often a long-time commitment with people choosing to feed birds for years^19^, and adjustments to feeding practices are more often in response to observations of nature than to human-centered constraints of time and money^12^. However, As scholarship on multispecies care has shown us^21^, this the care provided through bird-feeding is likely to be far from ‘innocent’ and can constitute a substantial power imbalance between participants^21^. We suggest that zooming into individual events and species appearing atin the bird feeder and then following reciprocal (re)actions should allows enable us to better understanding of humans’ relations to non-human species and vice versa. We hypothesize that any changes in feeding will practices expose important motivations, conflicts and trade-offs that people experience within between-species relations.

We utilise a large dataset of over 14,000 self-reported responses to an online questionnaire where residents of Finland were asked to describe events that have influenced changes in their bird-feeding practices over the last 20 years. As these changes relate to the actual events and their social and ecological consequences, we can delve deeper than inquiring about human behaviour in hypothetical situations. Bird-feeding has a long history in Finland spanning from the 19^th^ century ^20^ and currently bird-feeding is a widespread hobby across the country ^19^. Nevertheless, the number of people feeding is diminishing although the amount of food being provisioned has increased in some cases^19^. At the same time, bird communities are rapidly changing due to, e.g., climate change ^22,23^ with implications for broader ecological processes, particularly in urban environments^24^. Therefore, bird-feeding in Finland provides an ideal context to study human-bird interplay and multispecies relations. We asked (1) what explains the changes in people’s bird-feeding practices, and specifically, (2) which species affect bird-feeding practices. We expect that people will report reducing feeding activities when species perceived as unwanted appear at the feeding site, or the presence of wanted species will have led to increases in bird-feeding, with value-based statements thus revealing variation in attitudes towards different species.

## Materials and methods

### Questionnaire design and participation

We designed a questionnaire to better understand how and why people feed birds in Finland^19^. The questionnaire was prepared in Finnish and Swedish (two of the official languages of Finland) using the Google Forms platform and was available for two months between January 29th and March 31st, 2021. Participation was solicited openly through the social media channels of Finnish Museum of Natural History Luomus, through the local chapters of BirdLife Finland, and by publicity through television interviews and online news provided by the Finnish Broadcasting Company YLE, and the questionnaire could be accessed through the website of the Finnish Museum of Natural History Luomus or by direct link.

While the questionnaire included 13 questions^19^, only 4 questions with open-ended answers are analysed in detail here. It is noteworthy that the questionnaire did not limit respondents to answering about bird-feeding to e.g. only wintertime bird-feeding or to the feeding done in the respondents’ own yard, though these were clearly emphasized in the answers. The open-ended questions were (translated): “Have you changed your bird-feeding habits within the last approximately 20 years?”; “Why have you changed your feeding patterns?”; “If you have stopped feeding birds did any or several of the following factors influence the decision to quit?”; and “Give feedback to the creators of the questionnaire.” The first three of these questions had closed options, but also an open-ended alternative. The respondents that gave discrepant answers between closed and open alternatives (e.g., they had answered in closed option that they did not change feeding, but then explained in the open alternative answer the ways that they had changed feeding) were removed from further analysis.

### Qualitative analysis

We used data-driven content analysis to categorise individual responses through coding, i.e., “looking for pattern or order in a body of data by identifying recurring themes, categories or concepts” ^25^. The unit of analysis was either a full or partial answer which contained a comprehensible thought and these varied from single words to several sentences. Units of analysis were coded based on the direction of the change: decrease or increase. *Decrease* contained decrease in the amount of feed or the span of feeding time, moving the feeding site due to human-centric reasons (i.e., the feeding site was changed to make feeding easier for humans) or changing the feed to cheaper, whereas *increase* could be increase in the feed or the span of feeding time, moving the feeding site due to bird-related reasons (i.e., moving to a place with more birds or to safer location) and changing feed to a sort perceived more healthy or tastier. Codes also included the reason for the change whether this was a species or some other reason. Coding was done in qualitative data analysis software ATLAS.ti (v. 23.3.0 Mac; ATLAS.ti Scientific Software Development GmbH, Berlin, Germany). All coding and categorisations were completed using the original language of the respondent and translated into English (with assistance of native English speakers) for discussion with co-authors and presentation purposes.

Initial coding by JE generated a rich and diverse set of 834 codes to describe the whole phenomenon. Then the coding was focused by MA to take into account all instances when an organismal species was indicated as a reason for changing feeding practices. At this point, about 1 000 responses where cross-checked to assess the initial coding: the inter-rater reliability was 0.99. There were no mismatches between existing species names and responses, but additional species that were missing from the coding were added at this point increasing the set to 954 codes. Initial codes included also a number of answers which contained the respondents’ thoughts on why other people have changed their feeding practices and these were removed. Thus, the final set included 216 codes which were further classified thematically to create different categories of reasons and to include their directions of change.

Next, the focal taxa were selected. The initial codes contained sometimes long causal chains, such as nearby forest was clear-cut which led to decrease in the number of birds in the feeding site. In such cases, only the most proximal reason and taxon (here: lack of wanted birds) were used. Additionally, humans, the coronavirus, pet chickens and species offered as a feed, such as pork (lard) or peanut, were removed at this point.

The categorization of the reasons for the change in feeding was challenging: the number of codes was substantial and codes could create hierarchical relationships [e.g., (1) Corvids as (2) big birds spread food around, which (3) makes a mess, which in turn attracts (4) rats, which come (5) inside the building and do (6) damage]. The codes were therefore then organized and classified in visual networks. These networks were first built with the core codes that contained many responses, then organized into main categories and then further split into subcategories. Finally, the remaining codes were added one by one to the network. Each code belonged to only one subcategory, but the subcategories could include codes with opposite directions of change.

### Ethical considerations

The respondents gave informed consent for participation before entering their responses in the questionnaire. No personal data was asked for and none of the questions resulted in data collection did not fall under any of the categories warranting that institutional review would have been warranted under the rules of the Finnish National Board on Research Integrity (TENK). In open answers, the respondents could provide text give answers that could would be considered as identifiable personal or sensitive data (e.g., in explaining how a specific disease might prevent them from easily maintaining bird feeding site). These data were handled in line with European Union GDPR regulations and no answers that had identifiable or sensitive information were published.

## Findings

There were in total 15,088 open-ended answers from 14,215 respondents of whom 9,473 answered at least one open-ended question. As similar answers could be given to more than one open-ended question all answers by a single respondent have been analyzed together.

### Species and species groups affecting feeding practices

A total of 1,814 mentions of 58 different species or species groups were named as primary reasons for changing feeding practices (Table 1; Figure 1). Most of the species that affected bird-feeding habits were animals, and birds and mammals had the most mentions (85% of mentions; 49 out of 58 taxa). The most of mentioned taxa in all answers in general, as well as in only the answers linked to a negative change in feeding, were rats (586 mentions of decreased feeding, 1 of increased), followed by mice (269 decreased, 3 increased), corvids (156 decreased, 34 increased) and pathogens (111 decreased). Mammals were markedly related to decreased feeding, and they could even lead to stopping feeding altogether. In contrast, increased feeding is spread more evenly throughout the taxa. Waterfowl, thrushes and other medium-sized birds such as waxwings were the only ones that caused only increase.

**Table 1.** Classification of mentioned taxa within higher-level taxa. For each taxon, the columns show the number of mentions related to either decreased or increased feeding. Taxa are grouped to contrast the responses, wherever interesting; for example, woodpeckers more often led to decreased than increased feeding, whereas other medium-sized birds were without exception linked to increased feeding.

| Organismal group | Examples | Decrease | Increase |
| --- | --- | --- | --- |
| Pathogens | Bird flu, salmonella | 111 |  |
| Ectoparasites | Ticks, fleas | 7 |  |
| Waterfowls | Swans, ducks |  | 8 |
| Birds of prey | Owls and accipitrids | 6 | 2 |
| Other large-sized birds | Pigeons, seagulls, corvids | 187 | 46 |
| Woodpeckers | Black woodpecker, great spotted woodpecker | 11 | 8 |
| Other medium-sized birds | Thrushes, Bohemian waxwing |  | 20 |
| Tits | Great tit, blue tit | 10 | 18 |
| Other small birds | Sparrows, yellowhammer | 4 | 26 |
| Pets | Dog, cat | 40 | 36 |
| Wild carnivores | Raccoon dog, red fox, mustelids | 12 | 3 |
| Deer | White-tailed deer, roe deer | 7 | 13 |
| Hares | Hares, rabbit | 18 | 21 |
| Red squirrel |  | 99 | 67 |
| Other rodents | Brown rat, mice, voles | 1,005 | 5 |
| Shrews |  | 4 |  |
| Snakes |  | 1 |  |

**Figure 1:**
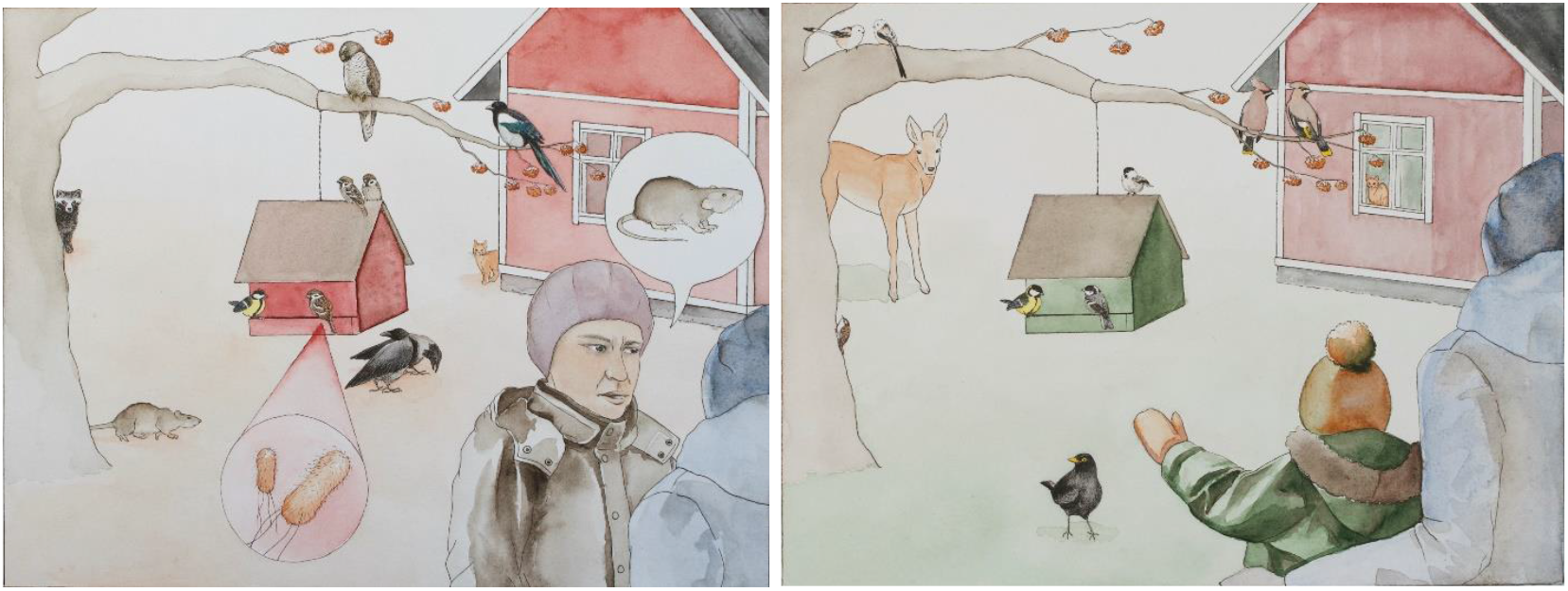
Graphical depiction of species and contexts that lead to decrease in feeding birds (left) and species and contexts that lead into increases in feeding birds (right). Drawn by Susu Rytteri.

Most of the mentioned taxa were linked both to increase and decrease in the feeding. For example, cats affected bird-feeding negatively when it was neighbours’ cats entering feeding site and attacking birds, whereas the increase was mostly due providing entertainment for own cats who followed the activities at the birdfeeder through window. It should be noted that positive or negative changes in feeding do not necessarily relate for person’s affective perception towards the species. For example, a species can eat so much food, that it requires increased food provision to make sure that also other species have enough food. This in turn could either lead to actual increase in the feeding or it could lead to a decrease if, for example, the person could not afford buying more food and decided to just stop feeding.

### Reasons for changing feeding

Based on the 4,992 responses which included a reason for change in bird-feeding, we formed three main categories for the reasons to change: 1) the respondent’s relation to another species, 2) the respondent’s relation to other humans and 3) relations between other species, and these were divided into seven subcategories and twenty-six individual reasons (Table 2). It is notable that on a main category level, both humans and non-humans can be present in any category. For example, the relations between humans are facilitated by complex chains of events:

**Table 2.** The main categories, subcategories and individual reasons for increasing or decreasing bird-feeding. The number of responses is shown on the level of subcategories and individual reasons. If reasons can be seen as symmetrical, i.e., the same reason can lead to both decrease and increase in the bird-feeding, those are placed on the same row in the table. The responses were given in Finnish or Swedish and translated here to English for presentation.

| Main category | Subcategory | n | Reason for increasing feeding | n | Example | Reason for decreasing feeding | n | Example |
| --- | --- | --- | --- | --- | --- | --- | --- | --- |
| Respondent's relation to non-human species | Attitude and interest towards nature or birds | 3,328 | Empathy, pity | 248 | "I noticed that the birds are very hungry when I gave them nuts from my hand." | Empathy, pity | 10 | "I do not think it is sensible to tinker with birds' habits and their probabilities in surviving the winter." |
|  |  |  | Interest in birds or nature | 306 | "I got interested in bird species identification." | Interest in birds or nature | 1 | "I moved the feeding in the shade so that it is easier to identify birds." |
|  |  |  | Healthier or tastier food to birds | 61 | "I've changed the feed that I provide based on what birds eat." | Picky birds | 17 | "Suet balls weren't eaten." |
|  |  |  | Supporting wanted species | 793 | "Because there's also other species than birds around, such as rabbits, deer, raccoon dogs and squirrels." | Lack of wanted birds or presence of unwanted birds | 1,892 | "Too many sparrows on the feeder, only two wanted species (great and blue tit) on the feeder." |
|  | Diseases | 941 |  |  |  | Zoonosis | 520 | "I have plants on the balcony and I don't want the birds to touch them as I'm worried about pathogens spreading from bird feces." |
|  |  |  |  |  |  | Pathogen spread between birds | 421 | "There were several dead birds two winters ago." |
|  | Economic reasons | 28 |  |  |  | Too much demand for food | 28 | "Jackdaws came to the feeder meant for the small birds. No food left for small birds and upkeep was expensive." |
|  | Scientific reasons | 6 | Doing research | 5 | "I do research on the overwintering of birds that use feeders." | Stopping doing research | 1 | "I used to ring birds in the winter." |
| Respondent's relation to humans | Social relations | 169 | Counterreaction to social pressure | 1 | "People are afraid of rats and blame people feeding. It just increases my interest in feeding birds and squirrels." | Social pressure | 91 | "My spouse prohibited [feeding] because of the rats." |
|  |  |  |  |  |  | Sabotage | 20 | "Neighbours are bothered, suet ball holders are stolen and [feeding] trees are being watched. All feeders are destroyed in the area." |
|  |  |  | Lack of human contact | 1 | "It is nice to observe birds especially due to social distancing (covid)." |  |  |  |
|  |  |  | Traditions | 30 | "I feel it is important to teach children how wildlife can be taken care of." | Children moving away | 11 | "Children have moved away. It was nice to observe birds with them." |
|  |  |  | Others are feeding | 1 | "I dared to start feeding as I saw my neighbor doing it." | Others are feeding | 14 | "Somewhere close by, there's better selection and birds are rarely in my yard. I do not want to get into an arms race." |
|  | Rules | 187 |  |  |  | Prohibitions | 187 | "The housing association has prohibited feeding due to an increase in the rat population." |
| Relations between non-human species | Ecological reasoning | 333 | Supporting biodiversity | 143 | "We want to support biodiversity around our cottage." | Effects on bird community | 69 | "Feeding changes the bird community too much." |
|  |  |  |  |  |  | Unnecessary | 22 | "I feed only from December to March as the birds do not need it otherwise." |
|  |  |  | Pets, domestic animals or garden | 24 | "Functions as a television for our indoor cats." | Pets, domestic animals or garden | 75 | "Tits come to the feeder and then they go to the barn to peck on cows' udders." |

> “My neighbours did not like the hulls of sunflower seeds on their terrace, so I changed to hulled seeds.”
>
> -Respondent 516

The primary reason for changing to more easily edible seeds in this case were the neighbours, i.e., other humans. At the same time, underlaying chain of events was that the neighbours were (perceived) to be annoyed by a plant part that was brought to their property by the birds. While arguably all chains of reasoning end up to the person feeding or not feeding birds, the important factor for the main category classification was where the respondents placed the primary reason for the change in their practices. Main category respondent’s relation to another species contained 86% of all classified responses and therein most were classified into the subcategory on the attitudes and interest towards birds and nature. The symmetric pair of disposition to feed certain species and not others *Supporting wanted species / Lack of wanted birds or presence of unwanted birds* were the most numerous and second most numerous reasons, respectively. In contrast, other people and relations between other species were both 7% of the reasons.

Many reasons formed symmetrical pairs, where the same event and reasoning could lead to different consequences on the feeding practices:

> “I pity birds during the freezing weather. “
>
> -Respondent 10706
>
> “[Birds] could have gotten used to feeding and, when there would have not been supplemental food, they would have suffered.”
>
> -Respondent 10

This could also relate to the events in the bird feeder itself, such as squirrels or magpies eating a lot could lead to either putting much more feed into the feeder or completely stopping the feeding. The decision depends on the economical issues, the effort needed and the respondent’s morales, if some species or individuals seemed, for example, to behave unfairly towards the other species or individuals. In general, the worry about biodiversity can relate to the survival of bird populations but also about how bird-feeding changes communities.

On main category level, the relation to humans led predominantly to negative change in feeding (33 positive changes to 323 negative), while the relation to another species and relations between non-human species were more evenly spread. Zoonosis and pathogen spread between birds, economic reasons, sabotage and (housing) rules were the reasons that led only to reduction in feeding. While other reasons had both directions, they were, though, usually clearly more negative or positive-oriented. For example, empathy and pity led mostly to increase (248 to 10) whereas social pressure led to decrease (1 to 91).

Ecological thinking also came into play to create more complex scenarios such as:

> “Too many mice, voles at the feeding site, leads to increase in snakes in the backyard.”
>
> – Respondent 8907

While rodents are commonly seen negatively, this respondent took a further step to link rodents to snake presence. Additionally, some were worried that birds of prey would eat smaller birds, whereas other found this as a reason to feed small birds to attract birds of prey to the backyard. This showed strong evidence of how people differentially value also species that are generally considered wanted species.

## Discussion

While the focus of bird-feeding studies has been either on the effects that bird-feeding has on avian or non-avian populations and communities^9,26–28^ or the reasons why humans feed birds^2,6,29–31^, our approach reveals a more complex diversity of ways in how the birds and other species attracted to bird-feeding sites can in turn affect how people provide food for birds. The species present or absent at the birdfeeder can both affect the bird-feeding practices, but, interestingly, most of the species can, depending on the context, lead to both increases or decreases in feeding birds. Context is highly important for many taxa: for example, cats are often seen as unwanted when a neighbor’s cat is predating birds at a feeding site, but when the owner’s cats are inside and watching birds through window, they can provide a reason to feed birds. Sometimes making decisions based on multispecies relations may lead to potential trade-offs, such as a respondent who said that there is no problem with the seeds that fall to the ground and attract rodents as freely-roaming cats kills those rodents. Indeed, our approach of asking about the changes in the feeding practices revealed a plethora of primary reasons that interact with species when people are making decisions on whether to put more or less effort into bird-feeding. Thus, clearly, bird-feeding is not a one-way interaction, but rather a series of reciprocal relations where humans and non-humans react to each other.

Our results show that the most common species that leads to a decrease in bird-feeding in Finland is the brown rat with a large majority of respondents describing this caused a reduction or even stopping of feeding birds. The most important reason to increase feeding was the occurrence of wanted species in the bird feeders. These are not surprising results as they correspond well to previous findings ^1,12,32,33^.

Nevertheless, closer inspection suggests more complex phenomena: the same taxa can not only lead to both an increase and decrease in different contexts, but also the same reason can lead to either an increase or a decrease of feeding. The red squirrel (*Sciurus vulgaris*) is an interesting example as it can be both a wanted or unwanted species at the bird feeder. Nevertheless, if a squirrel, regardless of its status, appears at the feeding site it could lead to an increase of provisioning to make sure that birds also get food, or it could lead to feeding birds being stopped altogether due to concern that the squirrel may eat too much food in comparison to what the respondent was prepared to invest in bird-feeding. Indeed, our results corroborate and expand on Dayer et al.’s ^12^ findings that there can be both positive and negative feedback loops between observations of species at the birdfeeders, human attitudes and actions, and subsequently changes in the species community at feeding sites. Similarly, the disappearance of a wanted species from the feeding site can lead to both increase and decrease in feeding birds. The respondents described ending the feeding completely, increasing the amount of feed or changing feed qualitatively in response to the situation. While some reasons can be expected to lead to a certain direction of change (for example, pathogens always lead to decreasing feeding), there is still a variation in whether the change is total stopping, reduction, moving the feeding site or changing the feed.

The respondents explained both straight-forward direct reasoning where a species was simply wanted and thus fed, whereas sometimes the reason was indirect and long and convoluted reasoning explained why a species was wanted or unwanted. For example, the long-tailed tit (*Aegithalos caudatus*) was considered to be cute and thus they were fed. In contrast, some people explained that they attract birds and ensure their survival to reduce the pest insect pressure for the garden in the summer. These findings are not surprising and they resonate with previous research: for example long-tailed tit is aesthetically pleasing to people ^34^. Some species, such as rats are a typical example of both direct and indirect reasonings as they can be unwanted based on just not liking them^35,36^, but also reasoned through zoonotic or infrastructure risks (even if the actual effect could be limited^37^). Indeed, these perceived risks can lead to regulations and prohibitions, which might lead to stopping feeding, even though the person feeding would have liked to continue (although our questionnaire exposed that regulations or social pressure are not always acted upon). Many respondents were also worried about the ecological effects of feeding birds. If bird-feeding has been initially seen as targeted at improving the welfare of individual birds^20^, nowadays respondents were clearly also thinking about the effects on specific, usually endangered species and also to the biodiversity at large. This can be seen, for example, in the responses where observing tits at the feeding site could also reduce feeding. There has been discussion about synanthropic tits (Eurasian blue tit, *Cyaniste caeruleus*; great tit; *Parus major*) benefitting at the expense of non-synanthropic tit species (willow tit, *Poecile montanus*; coal tit, *Periparus ater*): as their survival increases during the wintertime bird-feeding, they can dominate the community during the breeding season^38,39^. This knowledge has lead to some respondents reporting being worried about willow tits as a reason to stop bird-feeding altogether, although the recent results do not support this interaction between tit species^40^. Thus, different perceptions of the context could lead to different outcomes when it comes to feeding.

The most common relation that affected bird-feeding within our respondents was human-to-other species relation. This was expected as bird-feeding is generally expected to function on humans’ terms and conditions. For example, magpie can be seen as “stealing” food from the feeder: this might be due to people not wanting to feed magpies as they want to feed small birds or also because magpies, as large-sized birds, eat much more and while people might not have anything against magpies, they cannot afford feeding magpies. It seems that cultural norms such as small birds being in more apparent need of care than corvids are in play here. Nevertheless, other relations are also important. For example, social conventions, relations and norms affect the feeding. These norms can be explicit as written rules or neighbors shouting over the fence or implicit such as perceived acceptability in the neighborhood^29^. For example, urban apartment buildings are commonly administered as housing associations, where every apartment owner has a vote and these associations commonly prohibit bird-feeding in the common yard due to, e.g., rat presence^41^. Some reasoning could be also seen as caring for someone other than birds: many people care for non-human animal-to-animal relations, such as evidenced by providing entertainment for one’s pets.

Our findings should be understood within the Finnish context, especially when it comes to the quantitative results. For example, in Finland bird-feeding has long been more common during the winter, in particular when there is snow on the ground. Feeding birds when there is no snow on the ground is much less common and is more often directed to waterfowl or to birds in urban parks^19,42^. Although it is unknown how common bird feeding in Finland is, the total number of respondents to our questionnaire (i.e. more than 9,000) is close to 0.5 percent of all households in Finland (i.e., 2 793 636 in 2021)^43^. About half of the households in the UK feed birds^1^, but we expect this proportion to be lower in Finland. We have presented the frequencies of each of the answers, but we are not able to assess how representative these are of general population. While the answers were sometimes very short, they were quite informative, such as “My heart was so broken when the magpie found the suet balls.” We would expect that qualitatively our data would contain any important phenomena that would occur. We suggest that the underlaying phenomena might be quite universal, such as is shown by the global surge in bird-feeding during the pandemic^44^. On the non-human animal side, even if the particular species are different in different locations, their ecological niches vary similarly to Finland and are thus likely to present similar observations and interactions in bird-feeding sites. Thus, we suggest that while the specific species and proportions of different reasons vary from country to country, these relations are present across the globe.

Asking about the changes in the bird-feeding practices and their underlaying reasons directly links human actions and the relevant observations that humans make in their bird feeders. This approach does depend on the human perception about the change in the feeding. We do not know about the changes that are perceived as neutral nor can we be sure whether the humans’ perception of the direction of the change is the real effect on birds. For example, a study showed that while using a squirrel guard increased the amount of feed going to birds, it reduced bird visits to the bird feeder and had differential effects on different bird species^30^.

In conclusion, we suggest that the relationship between bird-feeding and humans’ animal attitudes is a highly dynamic process and also context-dependent: the same species can in different contexts lead to either increase or decrease of provisional feeding. Similarly, the same reasons can be used to justify increases and decreases in feeding. Thus, the avian and non-avian species that are directly or indirectly affected by the bird-feeding also affect the humans and become actionable participants in the process. This phenomenon has both ecological and social dimensions which are tightly interlinked. While bird-feeding influences species communities, it is also important to acknowledge and understand how other species influence feeding practices and behaviours and are active participants in bird-feeding practice.

## Acknowledgements

We thank Herre de Boendt, Risto Willamo and Perttu Seppä for their helpful and critical comments. This work was funded by Helsinki Institute of Sustainability grant to TA, Emil Aaltonen Foundation grant to TA, Maria de Maeztu Centre of Excellence IMEDEA (CSIC-UIB) (CEX2021-001198) to AS, European Commission Horizon 2020 Marie Skłodowska-Curie Actions (Grant no. 101027534) to AS and Helsinki Institute of Life Science grant to RT.

## Contribution

Research idea was formulated by MA, TA, AK, data collected by PD, AH, KR, AS, AL, RT, data analysed by MA, JE, HL, VV, AK, TA, the study was coordinated by TA and AK, while TA wrote the first draft of the manuscript while all authors reviewed and revised it.

## References

1. Cox, D. T. C. & Gaston, K. J. Human–nature interactions and the consequences and drivers of provisioning wildlife. Philosophical Transactions of the Royal Society B: Biological Sciences 373, (2018).

2. Cox, D. T. C. & Gaston, K. J. Urban Bird Feeding: Connecting People with Nature. PLoS One 11, e0158717 (2016).

3. Robb, G. N., McDonald, R. A., Chamberlain, D. E. & Bearhop, S. Food for thought: Supplementary feeding as a driver of ecological change in avian populations. Front Ecol Environ 6, 476–484 (2008).

4. Orros, M. E. & Fellowes, M. D. E. Wild Bird Feeding in an Urban Area: Intensity, Economics and Numbers of Individuals Supported. 10.3161/00016454AO2015.50.1.006 50, 43–58 (2015).

5. Shutt, J. D., Trivedi, U. H. & Nicholls, J. A. Faecal metabarcoding reveals pervasive long-distance impacts of garden bird feeding. Proceedings of the Royal Society B: Biological Sciences 288, (2021).

6. Reynolds, S. J., Galbraith, J. A., Smith, J. A. & Jones, D. N. Garden bird feeding: Insights and prospects from a north-south comparison of this global Urban phenomenon. Frontiers in Ecology and Evolution vol. 5 Preprint at 10.3389/fevo.2017.00024 (2017).

7. Reed, J. H. & Bonter, D. N. Supplementing non-target taxa: bird feeding alters the local distribution of mammals. Ecological Applications 28, 761–770 (2018).

8. Becker, D. J. & Hall, R. J. Too much of a good thing: Resource provisioning alters infectious disease dynamics in wildlife. Biol Lett 10, (2014).

9. Galbraith, J. A., Beggs, J. R., Jones, D. N. & Stanley, M. C. Supplementary feeding restructures urban bird communities. Proc Natl Acad Sci U S A 112, E2648–E2657 (2015).

10. Hanmer, H. J., Thomas, R. L. & Fellowes, M. D. E. Provision of supplementary food for wild birds may increase the risk of local nest predation. Ibis 159, 158–167 (2017).

11. Orros, M. E. & Fellowes, M. D. E. Supplementary feeding of wild birds indirectly affects the local abundance of arthropod prey. Basic Appl Ecol 13, 286–293 (2012).

12. Dayer, A. A. et al. Observations at backyard bird feeders influence the emotions and actions of people that feed birds. People and Nature 1, 1–14 (2019).

13. Clark, D. N., Jones, D. N. & James Reynolds, S. Exploring the motivations for garden bird feeding in south-east England. Ecology and Society 24, (2019).

14. Brock, M., Perino, G. & Sugden, R. The Warden Attitude: An Investigation of the Value of Interaction with Everyday Wildlife. Environ Resour Econ (Dordr) 67, 127–155 (2017).

15. White, M. E., Hamlin, I., Butler, C. W. & Richardson, M. The Joy of birds: the effect of rating for joy or counting garden bird species on wellbeing, anxiety, and nature connection. Urban Ecosyst 26, 755–765 (2023).

16. Cox, D. T. C. et al. Covariation in urban birds providing cultural services or disservices and people. Journal of Applied Ecology 55, 2308–2319 (2018).

17. Malone, K. Reconsidering Children’s Encounters With Nature and Place Using Posthumanism. Australian Journal of Environmental Education 32, 42–56 (2016).

18. Santangeli, A. et al. What drives our aesthetic attraction to birds? npj Biodiversity 2023 2:1 2, 1–7 (2023).

19. Deshpande, P. et al. How, why, where and when people feed birds? – Spatio-temporal changes in food provisionin in Finland. People and Nature in press, (2024).

20. Lähdesmäki, H., Aivelo, T. & Savolainen, P. Bird feeding devices exclude unwelcome visitors. Morethan-humans shaping the architecture and technology of birdfeeders in twentieth-century Finland. Environ Plan E Nat Space (2024) doi:10.1177/25148486241242680.

21. Puig de la Bellacasa, M. Matters of care: speculative ethics in more than human worlds. 265 (2017).

22. Antão, L. H. et al. Climate change reshuffles northern species within their niches. Nature Climate Change 2022 12:6 12, 587–592 (2022).

23. Hällfors, M. H. et al. Recent range shifts of moths, butterflies, and birds are driven by the breadth of their climatic niche. Evol Lett 8, 89–100 (2024).

24. Deshpande, P., Johansson, N., Kluen, E., Lehikoinen, A. & Thorogood, R. Changing bird migration patterns have potential to enhance dispersal of alien plants from urban centres. Glob Chang Biol in press,.

25. MacLure, M. Chapter 9 Classification or Wonder? Coding as an Analytic Practice in Qualitative Research. Deleuze and Research Methodologies 164–183 (2022) doi:10.1515/9780748644124-011/HTML.

26. Wilcoxen, T. E. et al. Effects of bird-feeding activities on the health of wild birds. Conserv Physiol 3, (2015).

27. Plummer, K. E., Risely, K., Toms, M. P. & Siriwardena, G. M. The composition of British bird communities is associated with long-term garden bird feeding. Nat Commun 10, (2019).

28. Robb, G. N. et al. Winter feeding of birds increases productivity in the subsequent breeding season. Biol Lett 4, 220–223 (2008).

29. Baverstock, S., Weston, M. A. & Miller, K. K. A global paucity of wild bird feeding policy. Science of the Total Environment 653, 105–111 (2019).

30. Hanmer, H. J., Thomas, R. L. & Fellowes, M. D. E. Introduced Grey Squirrels subvert supplementary feeding of suburban wild birds. Landsc Urban Plan 177, 10–18 (2018).

31. Jones, D. N. & James Reynolds, S. Feeding birds in our towns and cities: A global research opportunity. J Avian Biol 39, 265–271 (2008).

32. George, K. A., Slagle, K. M., Wilson, R. S., Moeller, S. J. & Bruskotter, J. T. Changes in attitudes toward animals in the United States from 1978 to 2014. Biol Conserv 201, 237–242 (2016).

33. German, D. & Latkin, C. A. Exposure to urban rats as a community stressor among low-income urban residents. J Community Psychol 44, 249–262 (2016).

34. Haukka, A., Lehikoinen, A., Mammola, S., Morris, W. & Santangeli, A. The iratebirds Citizen Science Project: a Dataset on Birds’ Visual Aesthetic Attractiveness to Humans. Scientific Data 2023 10:1 10, 1–12 (2023).

35. Bjerke, T. & Østdahl, T. Animal-related attitudes and activities in an urban population. Anthrozoos 17, 109–129 (2004).

36. Kellert, S. R. American attitudes toward and knowledge of animals: An update. in Advances in animal welfare science 1984 (eds. Fox, M. W. & Mickley, L. D. 177–213 (Springer, Dordrecht, Netherlands, 1985).

37. Arrindell, W. A. Phobic dimensions: IV. The structure of animal fears. Behaviour Research and Therapy 38, 509–530 (2000).

38. Orell, M. Population fluctuations and survival of Great Tits Par us major dependent on food supplied by man in winter. Ibis 131, 112–127 (1989).

39. Sofaer, H. R., Flather, C. H., Jarnevich, C. S., Davis, K. P. & Pejchar, L. Human-associated species dominate passerine communities across the United States. Global Ecology and Biogeography 29, 885–895 (2020).

40. Lehikoinen, A. et al. Population collapse of a common forest passerine in northern Europe as a consequence of habitat loss and decreased adult survival. For Ecol Manage 572, 122283 (2024).

41. Nygren, N. V & Tuomas, J. A. Rotta kuntalaisena – rottien esiintyminen ja hallinta [Rat as a municipal citizen – rat occurrence and control]. Suomen Eläinlääkärilehti 128, 331–337 (2022).

42. Haapanen, E. Villisorsista pullasorsiksi: eläinten kotiutumisesta kaupunkiin. in Näkökulmia Helsingin ympäristöhistoriaan, Kaupunki ja sen ympäristö 1800-ja 1900-luvulla. (eds. Laakkonen, S., Laurila, S., Kansanen, P. & Schulman, H.) 110–123 (Edita/Helsingin kaupungin tietokeskus, Helsinki, 2001).

43. Official Statistics of Finland (OSF): Dwellings and Housing Conditions. Reference Period: 2024, 2nd Quarter. https://stat.fi/en/publication/clm6ays13jvcx0buhrnat7zfb.

44. Doremus, J., Li, L. & Jones, D. Covid-related surge in global wild bird feeding: Implications for biodiversity and human-nature interaction. PLoS One 18, (2023).

